# Discordant inheritance of chromosomal and extrachromosomal DNA elements contributes to dynamic disease evolution in glioblastoma

**DOI:** 10.1101/081158

**Authors:** Ana C. deCarvalho, Hoon Kim, Laila M. Poisson, Mary E. Winn, Claudius Mueller, David Cherba, Julie Koeman, Sahil Seth, Alexei Protopopov, Michelle Felicella, Siyuan Zheng, Asha Multani, Yongying Jiang, Jianhua Zhang, Do-Hyun Nam, Emanuel F. Petricoin, Lynda Chin, Tom Mikkelsen, Roel G.W. Verhaak

## Abstract

To understand how genomic heterogeneity of glioblastoma (GBM) contributes to the poor response to therapy, which is characteristic of this disease, we performed DNA and RNA sequencing on GBM tumor samples and the neurospheres and orthotopic xenograft models derived from them. We used the resulting data set to show that somatic driver alterations including single nucleotide variants, focal DNA alterations, and oncogene amplification in extrachromosomal DNA (ecDNA) elements were in majority propagated from tumor to model systems. In several instances, ecDNAs and chromosomal alterations demonstrated divergent inheritance patterns and clonal selection dynamics during cell culture and xenografting. Longitudinal patient tumor profiling showed that oncogenic ecDNAs are frequently retained after disease recurrence. Our analysis shows that extrachromosomal elements increase the genomic heterogeneity during tumor evolution of glioblastoma, independent of chromosomal DNA alterations.

## INTRODUCTION

Cancer genomes are subject to continuous mutagenic processes in combination with an insufficient DNA damage repair ^1^. Somatic genomic variants that are acquired prior to and throughout tumorigenesis may provide cancer cells with a competitive advantage over their neighboring cells in the context of a nutrition- and oxygen-poor microenvironment, resulting in increased survival and/or proliferation rates ^2^. The Darwinian evolutionary process results in intratumoral heterogeneity in which single cancer-cell-derived tumor subclones are characterized by unique somatic alterations ^3^. Chemotherapy and ionizing radiation may enhance intratumoral evolution by eliminating cells lacking the ability to deal with increased levels of genotoxic stress, while targeted therapy may favor subclones in which the targeted vulnerability is absent ^4,5^. Increased clonal heterogeneity has been associated with tumor progression and mortality ^6^.

Computational methods that analyze the allelic fraction of somatic variants identified from high throughput sequencing data sets are able to infer clonal population structures and provide insights into the level of intratumoral clonal variance ^7^. Glioblastoma (GBM), a WHO grade IV astrocytoma, is the most prevalent and aggressive primary central nervous system tumor. GBM is characterized by poor response to standard post-resection radiation and cytotoxic therapy, resulting in dismal prognosis with a 2 year survival rate around 15% ^8^. The genomic and transcriptomic landscape of GBM has been extensively described ^9–11^. Intratumoral heterogeneity in GBM has been well characterized, in particular with respect to somatic alterations affecting receptor tyrosine kinases ^12–14^. To evaluate how genomically heterogeneous tumor cell populations are affected by selective pressures arising from the transitions from tumor to culture to xenograft, we performed a comprehensive genomic and transcriptomic analysis of thirteen GBMs, the glioma-neurosphere forming cultures (GSC) derived from them, and orthotopic xenograft models (PDX) established from early passage neurospheres. Our results highlight the evolutionary process of GBM cells, placing emphasis on the diverging dynamics of chromosomal DNA alterations and extrachromosomally amplified DNA elements in tumor evolution.

## RESULTS

### Genomic profiling of glioblastoma, derived neurosphere and PDX samples

We established neurosphere cultures from 12 newly diagnosed and one matched recurrent GBM (Table 1). Neurosphere cultures between 7 and 18 passages were used for molecular profiling and engrafting orthotopically into nude mice. The sample cohort included one pair of primary (HF3016) and matching recurrent (HF3177) GBM. A schematic overview of our study design is presented in Fig. 1a. To determine whether model systems capture the somatic alterations that are thought to drive gliomagenesis, and whether there is selection for specific driver genes, we performed whole genome sequencing at a median depth of 6.5X to determine genome wide DNA copy number as well as exome sequencing on all samples. DNA copy number was generally highly preserved between tumor and derived model systems (Supplementary Fig. 1). Whole chromosome 7 gain and chromosome 10 loss were retained in model systems when detected in the tumor, consistent with their proposed role as canonical GBM lesions that occur amongst the earliest events in gliomagenesis ^15^. The global DNA copy number resemblance between xenografts and the GBM from which they were derived confirms that PDXs recapitulate the majority of molecular properties found in the original tumor. We compared mutation and DNA copy number status of genes previously found to be significantly mutated, gained, or lost in GBM ^9,11^. We found that 100% of homozygous deletions and somatic single nucleotide variants (sSNVs) affecting GBM driver genes in tumor samples were propagated to the neurospheres and xenografts, including non-coding variants in the *TERT* promoter (Fig. 1b). Genomic amplifications showed greater heterogeneity. In two cases, *MYC* amplification was not detected in the parental tumor, but presented in the derivative neurospheres and maintained in xenografts, consistent with its role in glioma stem cell maintenance ^16,17^. Other genes showing variable representation across tumor and model systems included *MET* in *HF3035* and *HF3077*, and *EGFR* and *PIK3CA* in HF2354. The HF2354 derived model systems were considerably less similar compared to the primary tumor than other cases which coincided with HF2354 being the only case subjected to neoadjuvant carmustine treatment. Whole chromosome gains of chromosome 1, 14 and 21, and one copy loss of chromosome 3, 8, 13, 15 and 18 were acquired in the neurosphere culture and propagated to the xenograft models (Supplementary Fig. 1). At the gene level, this resulted in newly detected mutations in *PTEN* and *TP53*, focal amplification of *MYC* (also in HF3016), and absence of *CDK4* and *EGFR* amplification in the neurosphere and xenografts relative to the tumor sample (Fig. 1b).

**Figure 1.**
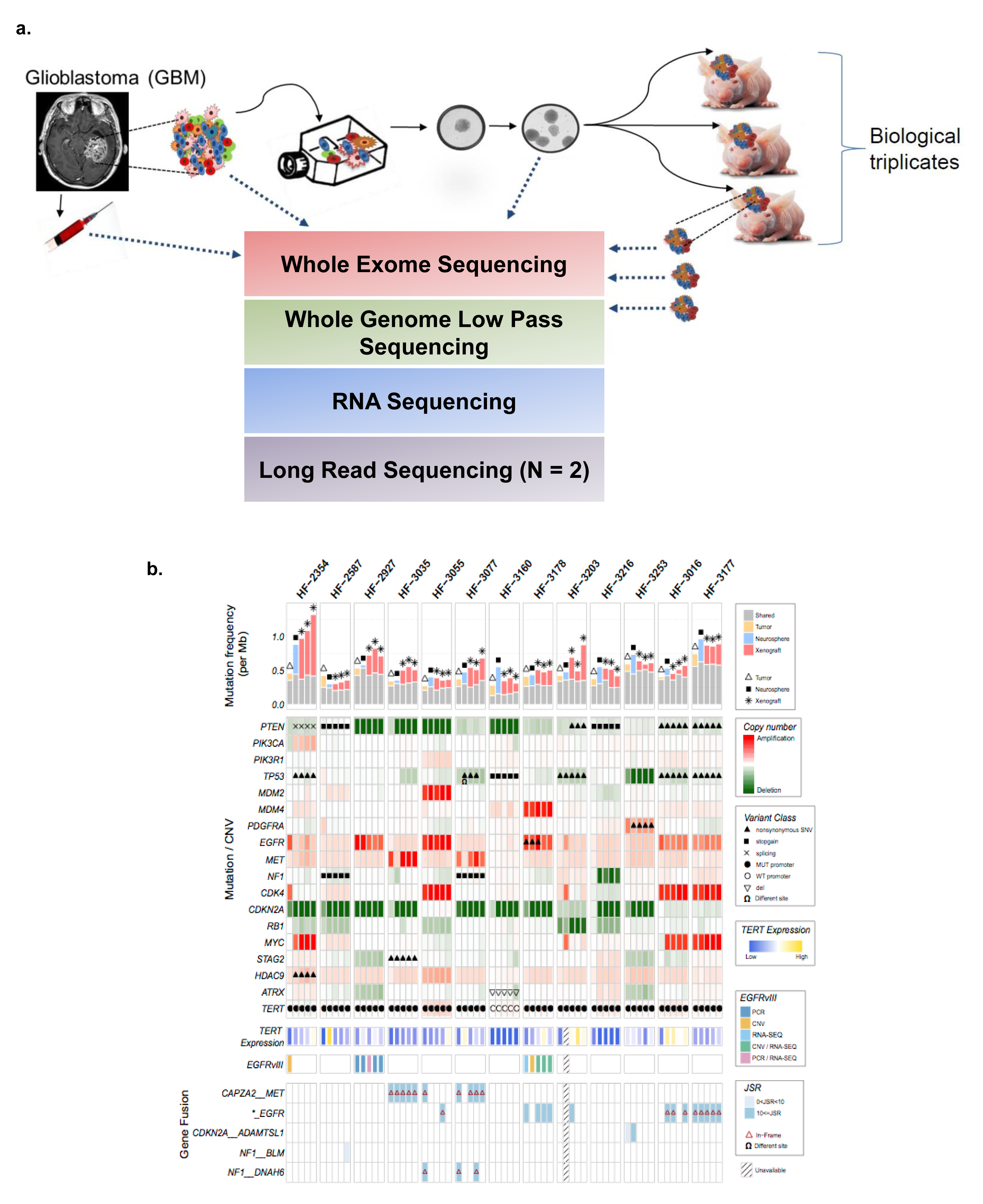
Comprehensive comparison of GBM, derived neurospheres and PDX models. **a.** Schematic study overview. **b.** Somatic driver alterations compared between GBM tumors and derivative model systems.

**Table 1.** Clinical characteristics of GBM patients included in this study.

| Sample | Pathology | Age/<br>Gender | Rx<br>prior to surgery | MGMT | OS<br>(days) | TTP<br>(days) |
| --- | --- | --- | --- | --- | --- | --- |
| HF2354 | GBM | 61/M | BCNU | U | 196 | 60 |
| HF2587 | GBM | 56/F | untreated | M | 360 | 232 |
| HF2927 | GBM | 55/F | untreated | U | 664 | 566 |
| HF3016 | GBM | 45/M | untreated | U | 649 | 88 |
| HF3177 | rGBM4 |  | RT/TMZ/DCVax | U |  |  |
| HF3035 | GBM | 54/F | untreated | U | 352 | 196 |
| HF3055 | GBM | 58/M | untreated | U | 371 | 77 |
| HF3077 | GBM | 56/F | untreated | U | 465 | 54 |
| HF3160 | GBM | 21/F | untreated | M | 1018 | 100 |
| HF3178 | GBM | 65/M | untreated | U | 189 | 138 |
| HF3203 | GBM | 64/M | untreated | U | 425 | 276 |
| HF3216 | GBM | 76/M | untreated | U | 94 |  |
| HF3253 | GBM | 82/F | untreated | U | 68 |  |

Rx: treatment; MGMT: *MGMT* gene promoter methylation status, U = unmethylated, M = methylated; OS: overall survival; TTP: time to progression.

### Extrachromosomal elements are frequently found in glioblastoma

Cytogeneticists have since long recognized that DNA in cancer can be amplified as part of chromosomal homogenously staining regions (HSR) and as extrachromomal minute bodies ^18^. An early example of the importance of extrachromosomal DNA elements (ecDNA) in cancer was the discovery of double minutes carrying the oncogene *N-MYC* in neuroblastoma ^19^. A recent survey of a compendium of cancer cells and cell lines highlighted the frequent presence of ecDNA in glioblastoma, among other cancer types,^20^, confirming previous studies ^21–23^. We searched our data set for complex patterns of DNA copy number amplification and rearrangement that are suggestive of ecDNA elements (Supplementary Fig. 2). On the basis of DNA copy number patterns we predicted 74 ecDNAs originating from 21 unique genomic loci which were distributed over ten of the thirteen patient tumors and their derived model systems. The predicted ecDNA elements contained oncogenes including *MYC*, *MYCN*, *EGFR*, *PDGFRA*, *MET,* the *MECOM/PIK3CA/SOX2* gene cluster and the *CDK4/MDM2* gene cluster. In total, 19 of the 21 unique oncogene carrying ecDNAs were detected in more than one sample, i.e. in neurospheres and matching PDX or in tumor sample and matching neurosphere or PDX (Fig. 2a). We performed interphase FISH on tumor samples and PDX, and metaphase FISH on neurospheres to validate 34 predicted ecDNA amplifications, including of *EGFR* (HF2927, HF3178, HF3016 and HF3177), *MYC* (HF2354, HF3016 and HF3177), *CDK4* (HF3055, HF3016 and HF3177), *MET* (HF3035 and HF3077), *MDM2* (HF3055) and *PDGFRA* (HF3253). In all interphase FISH experiments we observed a highly variable number of fluorescent signals per nucleus, ranging from two to 100 (Fig. 2b, Supplementary Table 1). This heterogeneity was strongly suggestive of differences in the number DNA copies of the targeted gene per cell and thereby of an extrachromosomal DNA amplification. Metaphase FISH on neurosphere cells validated the extrachromosomal status in all cases (Fig. 2b). Our analysis showed that oncogene amplification frequently resided on extrachromosomal DNA elements.

**Figure 2.**
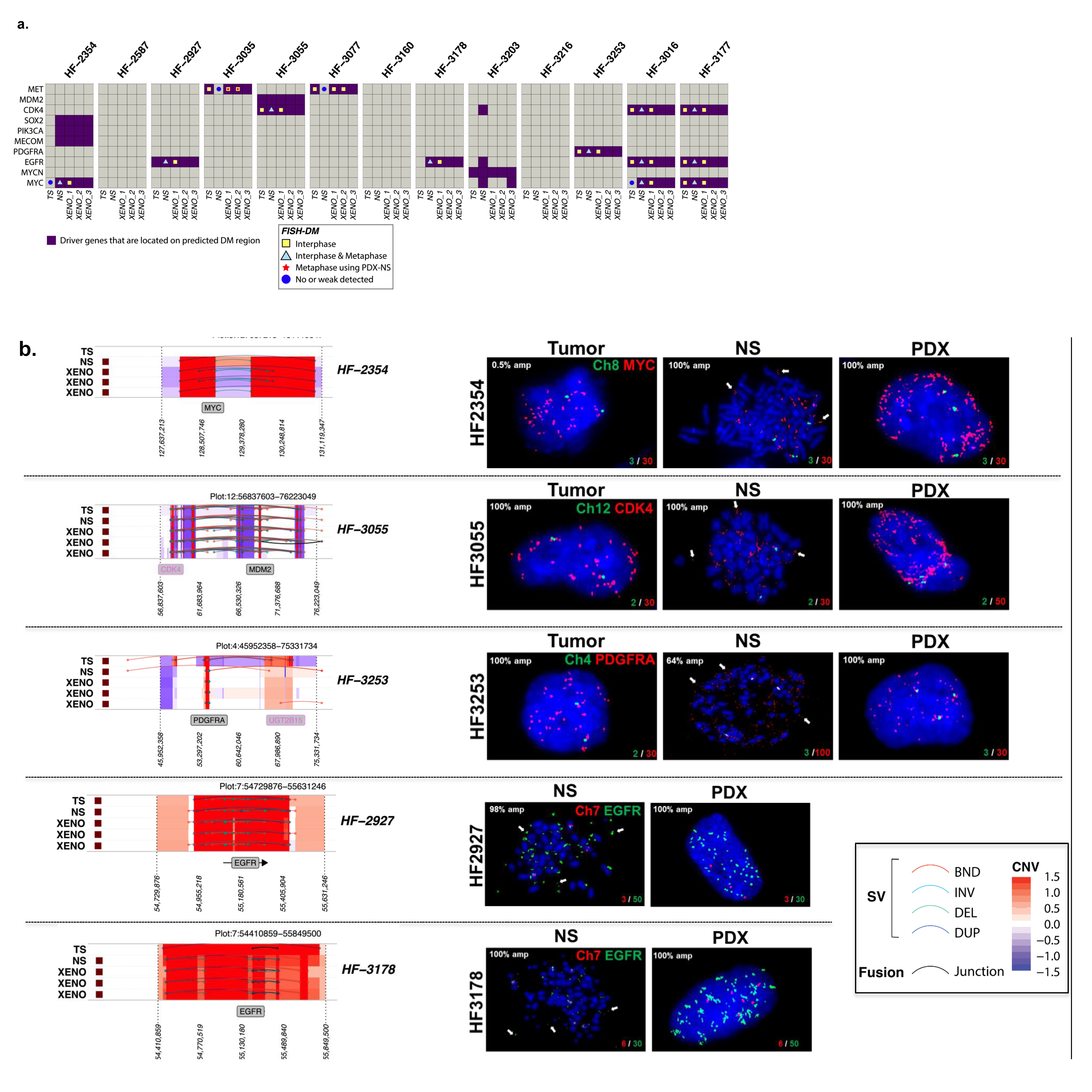
ecDNA in hGBM samples and FISH validation. **a.** Driver genes located on the potential double minute (DM) regions. **b.** Representative FISH images showing amplification of *MYC, CDK4, PDGFRA* in tumor, neurospheres and PDXs (red) and control chromosomal probes (green). *EGFR* amplification in neurospheres and PDX (green) and Chr7 control are shown. Metaphase FISH is shown for the neurospheres, with arrows pointing to extrachromosomal amplification.

### Extrachromosomal *MET* DNA elements mark a distinct tumor subclone

Among the identified oncogene carrying ecDNA elements, two cases of extrachromosomal *MET* amplification stood out due to their variable presence across the parental tumor (high frequency), neurosphere (low frequency) and xenograft triplicates (high frequency) (Fig. 3a). In both cases, the *MET* amplification associated with a transcript fusion with neighboring gene *CAPZA2* (Fig. 3b, Supplementary Fig. 3a). The pattern of undetectable and re-appearing *MET* rearrangements may result from clonal selection of glioblastoma cells with a competitive advantage for proliferation *in vivo*. This hypothesis is strengthened by the observation that the breakpoints of the lesions were identical across samples from the same parental origin (Supplementary Fig. 3b). MET is a growth factor responsive cell surface receptor tyrosine kinase and may provide context dependent proliferative signals ^24^. We reasoned that evolutionary patterns resulting in such dominant clonal selection would likely be replicated by sSNVs tracing the cells carrying the MET amplicon. To evaluate clonal selection patterns, we determined variant allele fractions of all sSNVs identified across HF3035 and HF3077 samples. To increase our sensitivity to detect mutations present in small numbers of cells, we corroborated the exome sequencing data using high coverage (>1,400x) targeted sequencing. All mutations detected in the HF3035 GBM were recovered in the neurosphere and xenografts. The mutational profile of HF3035 suggested that a subclone developed in the xenografts that was not present in parental GBM and neurosphere and revealed a subclone that was present at similar frequencies in all samples (Fig. 3c). Only a single and very low frequency *LAMB1* mutation (variant allele fraction in tumor = 0.003) present in the HF3077 primary tumor, but not detected in its derived neurosphere, resurfaced in one of three xenografts with a 0.04 variant allele fraction. A low frequency subclone (C2) developed in the neurosphere which was transmitted to xenografts (Fig. 3c). Subclonal heterogeneity as recovered by the mutation profiles thus suggested a very different clonal selection trend compared to to the disappearing and resurfacing *MET* amplifications and associated transcript fusions. EcDNAs are thought to inherit through random distribution over the two daughter cells^25^, possibly through a binomial model^26^, but much is unknown with respect to the propagation of ecDNA through cancer cell populations. The disjointed propagation of chromosomal SNVs and extrachromosomal *MET* ecDNAs indicate that they are marking different tumor subclones and suggest alternative modes of tumor evolution. While sSNVs are copied to daughter cells during mitosis such that both cells inherit the full spectrum of chromosomal alterations present in the parental cell, ecDNA elements likely randomly segregate and end up in the daughter cells in uneven numbers.

**Figure 3.**
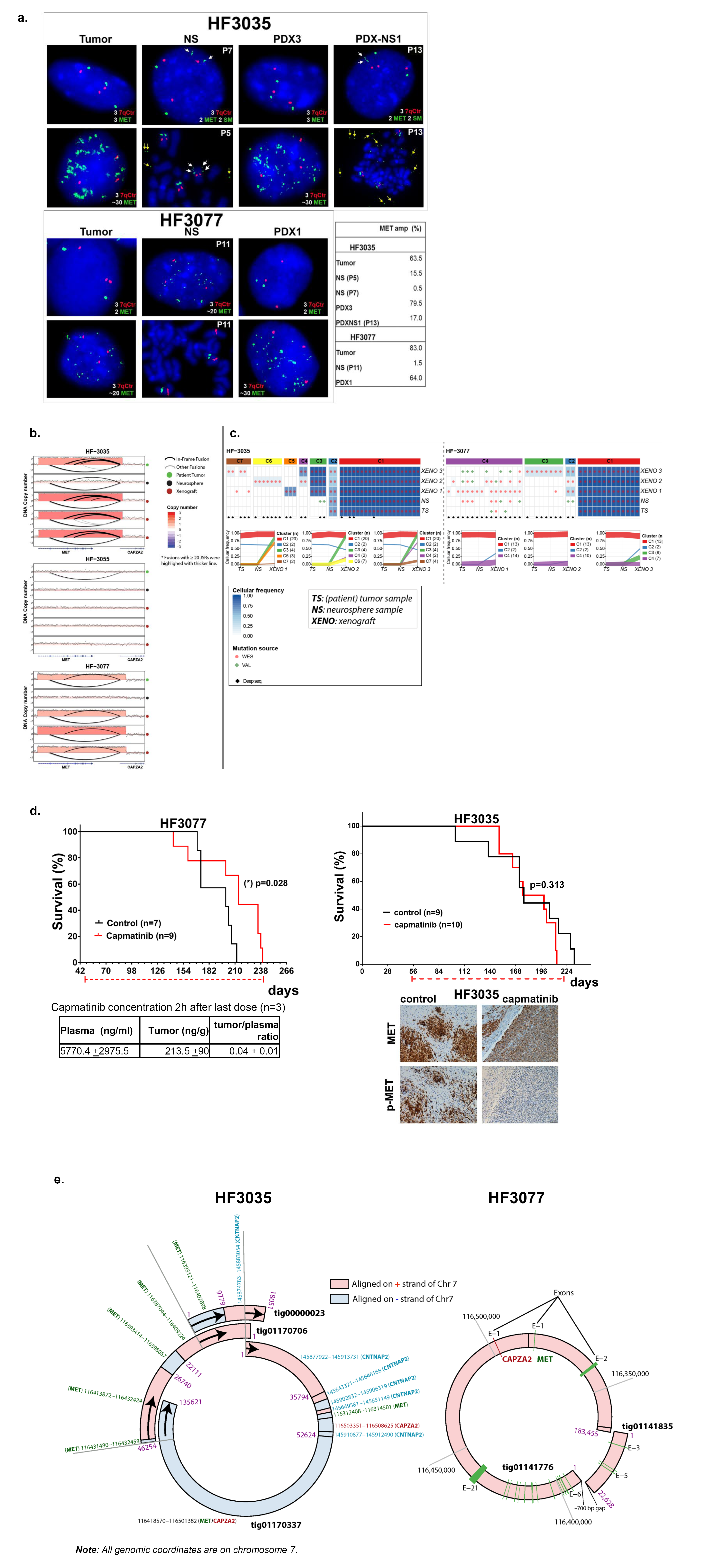
Extrachromosomal *MET* DNA. **a.** Representative fluorescent in-situ hybridization images for MET (green) and chromosome 7 control probes (7qCtr, red) labeling of HF3035 and HF3077 tumor, neurosphere (NS), and xenografts (PDX), and neurospheres established from HF3035 xenograft tumors (PDX-NS1). Passage numbers are indicated for neurosphere cultures. White arrows point to 2 fragmented MET signals in one chromosome in HF3035 samples (2SM). Yellow arrows point to double minute MET in metaphase nuclei of HF3035 neurospheres. The percentage of nuclei presenting MET amplification for each sample is shown. **b.** The 7q31 locus in three sets of GBM tumors and derivate models. **c.** Coverage-controlled sSNVs detected using exome and deep sequencing (top panel). Color reflects cellular frequency estimates. Bottom panel shows clonal tracing from HF3035 and HF3077 parent tumor to neurospheres and PDXs. Each line represents a group of mutations computationally inferred to reflect a subclone. **d.** Treatment with single agent capmatinib (30 mg/kg, daily oral doses) increases survival of HF3077 PDX, but not of HF3035. Kaplan-Meier survival curves were compared by log-rank (Mantel-Cox) test, significance set at P<0.05 (*), treatment schedule (doted red line) and number of mice in each arm (n) are shown. Capmatinib concentration in the plasma and tumor tissue collected 2h after the last dose was determined by LC-MS/MS for HF3077 PDX. MET and p-MET detection by IHC of control and capmatinib-treated xenografts show complete inhibition of p-MET, but did not affect MET overexpression in HF3035 PDX. Scale, 40 μm. **e.** Double minute structures predicted with long read sequencing in HF3035 and HF3077 xenografts.

MET expressing cells exhibited MET activation and were selected early during tumor formation in the orthotopic xenografts (Supplementary Fig. 3c), suggesting that MET activity was driving selection for *MET* amplified cells in vivo. Treatment of HF3077 PDX with ATP-competitive MET inhibitor capmatinib (INCB28060) ^27^ at a daily oral dose of 30 mg/kg showed a significant survival benefit, despite the relatively low concentration of drug in the brain tumor as assessed by LC-MS/MS (Fig. 3d). In contrast, capmatinib treatment of HF3035 PDX did not increase survival nor decrease MET expression but resulted in decrease of phospho-MET in treated tumors. This may reflect MET functions that are independent of the kinase activity in these tumors, as previously proposed ^28,29^. These results demonstrate that targeting MET in GBM harboring *MET* ecDNA amplification has therapeutic potential, but MET amplification alone is not a predictor of response to single agent ATP-competitive inhibitor treatment. Comparable to the orthotopic xenografts, subcutaneous PDX tumors formed from implant of HF3035 neurosphere cells were dominated by *MET*-amplified cells accompanied by robust MET expression (Supplementary Fig. 3c). The increase in the frequency of MET-amplification in HF3035 cells in vivo are therefore not dependent on factors uniquely present in the brain microenvironment.

Different genetic origins for ecDNA have been postulated, with evidence for post-replicative excision of chromosomal fragments and non-homologous end joining ^30^. Interphase FISH analysis in the parental HF3077 tumor identified a small percentage of nuclei with 3 copies of chromosome 7 but only 2 copies of *MET*. The frequency of cells with one deleted copy of *MET* in Ch 7 increased significantly in HF3077 neurospheres and decreased in the xenografts (Supplementary Table 1). The observed gene deletion in one copy of chromosome 7 is suggestive of the post-replication segregation-based model of double minute formation^30^. To precisely define the genomic contents and structure of the predicted double minutes, we generated long read (Pacific Biosciences) DNA sequencing from a single xenograft of each HF3035 and HF3077, and performed *de novo* assembly. In HF3035, seven assembled contigs (range: 6,466 ∼ 135,621 bp) were identified to have sequence fragments (at least 1,000 bp long) aligned on the *MET-CAPZA2* region of hg19 chromosome 7. Interestingly, analysis of the aligned sequence fragments from the seven contigs revealed a more complex structural rearrangement than expected from the analysis of short read sequencing data. For example, the 135kb tig01170337 contig consisted of 8 sequence framents that were nonlinearly aligned on alternating strands of the *MET-CAPZA2* and *CNTNAP2* regions. Other contigs such as tig01170699, tig01170325, and tig00000023 also showed nonlinear alignment, suggesting that these contigs resulted from chromosomal structural variations. We performed pairwise sequence comparison of the contigs to search for sequence fragments (at least 5,000 bp long) shared among them, and we found four contigs each of which shared sequence fragments with one of the contigs. Interestingly, three of them could be connected in a circular form using the shared sequence fragments (Fig. 3e; Supplementary Fig. 4a), revealing a circular structure that may represent the full double minute. In HF3077, only two contigs were detected to be aligned on the *MET-CAPZA2* region of hg19 chromosome 7 (Fig. 3e; Supplementary Fig. 4a). Presence of only two aligned contigs in HF3077 might be related to the lower sequence coverage of the double minute structure, compared to HF3035 (34x vs 405x, respectively) (Supplementary Fig. 4b). The longest contig, tig01141776 (183,455 bp long), consisted of two segment framents that were nonlinearly aligned over exon 1 of *CAPZA2* and all except exons 3-5 of *MET*, suggesting that it resulted from structural variations. The second short contig, tig01141835 (22,628 bp long), was aligned as a whole over exon 3-5 of *MET*. Interestingly, connecting the two contigs created a circular DNA segment. Through analysis of PacBio sequencing, we were able to detect and reconstruct the predicted double minute structures.

**Figure 4.**
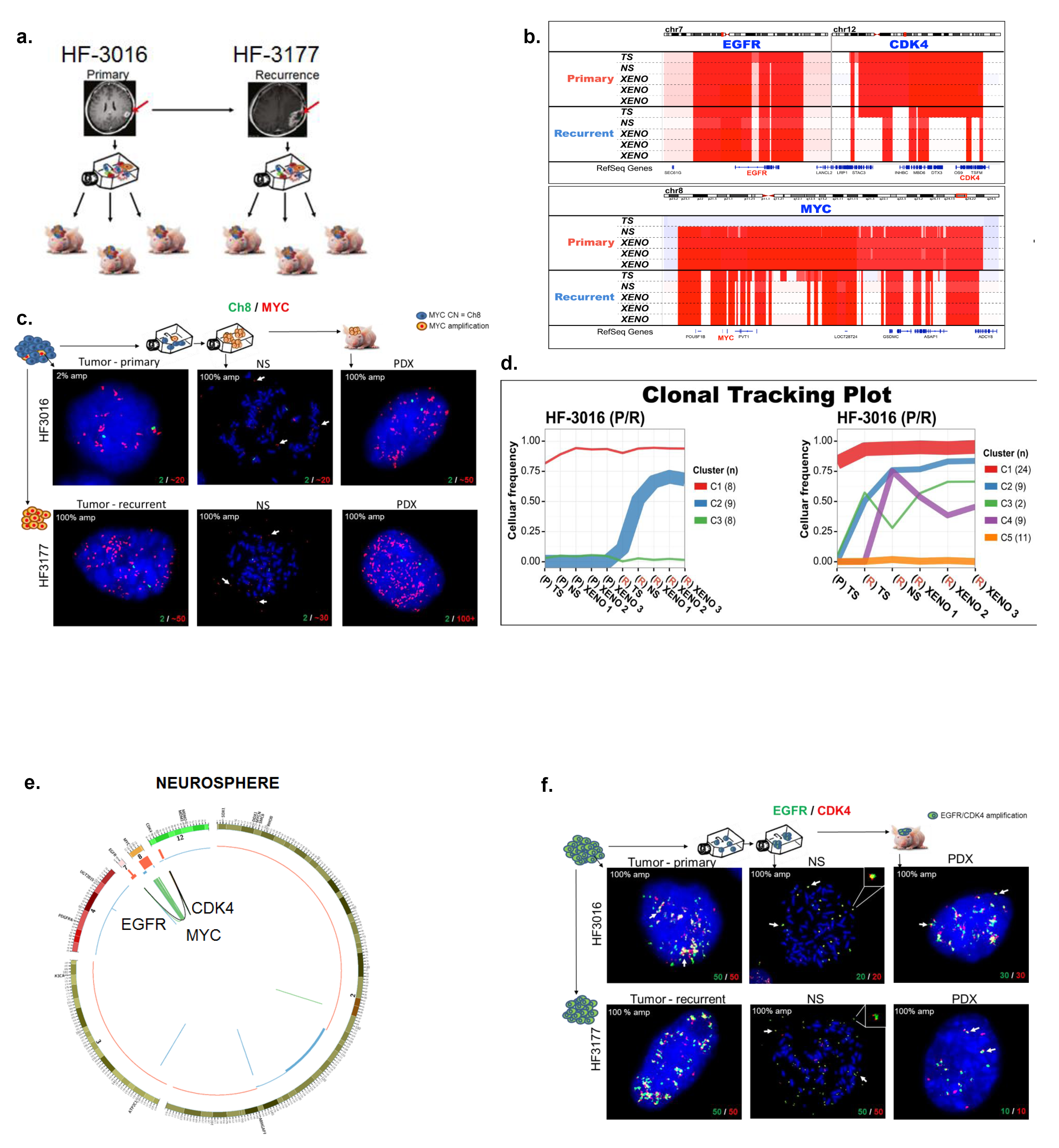
Double minutes drive tumor progression in patient tumors and derived model systems. **a.** Establishing neurosphere cultures and PDX models from a paired primary/recurrent GBM. **b.** A co-amplification of *EGFR* (chr7)/*CDK4* (chr 12) is detected in primary GBM HF3016 and this co-amplification is sustained in both neurosphere and xenografts derived from this primary tumor, as well as the recurrent GBM HF3177, and the neurosphere/xenografts thereof. The HF3016 primary tumor is not MYC amplified. The HF3016 neurosphere, as well as all HF3177 samples, show focal *MYC* amplification. **c.** Representative FISH images for MYC (red) and Ch8 marker (green) show that a small fraction (2%) of the cells in HF3016 tumor presents MYC amplification, while 100% of nuclei in the remaining samples present MYC amplification, which is clearly extrachromosomal (white arrows) in the metaphase spreads (NS). **d.** Clonal tracing of a pair of primary-recurrent GBM, their matching neurospheres, and xenografts. **e.** Starting in the neurosphere of the primary tumor, a complex structural variant is identified that connects the *CDK4* locus to the *EGFR* locus. The *MYC* locus is not part of this variant. The EGFR/CDK4 variant is detected in HF3016 PDXs as well as all HF3177 samples. f. EGFR (green) and CDK4 (red), detected by FISH, are amplified in 100% of nuclei for every sample from this patient, with identical copy numbers in each nucleus (bottom of the panels). Overlapping dots show that EGFR/CDK4 co-localize (white arrows) and metaphase FISH (NS) shows extra chromosomal co-amplification in the same double minute (inserts)

### Multiple ecDNA elements are longitudinally preserved in a patient GBM and its derivative model systems

Analysis of a pair of primary and recurrent GBM included in our cohort, respectively HF3016 and HF3177, showed that chromosomal and extrachromosomal elements jointly orchestrated complex evolutionary dynamics (Fig. 4a). Primary and recurrent tumor were globally very similar (Fig. 1b, Supplementary Fig. 1). While the HF3016 primary tumor showed diploid *MYC* DNA copy numbers, a focal *MYC* amplification was detected in the neurosphere and PDXs derived from this tumor, and the same *MYC* amplification was identified in all samples from the recurrent tumor (Fig. 4b). Interestingly, FISH analysis showed that *MYC* amplification was present in low frequency (2%) in the initial HF3016 tumor, and was enriched to 100% of nuclei in the neurospheres and in the recurrent tumor (Fig. 4c, Supplementary Table 1). Metaphase FISH analysis confirmed extrachromosomal *MYC* amplification in both HF3016 and HF3177 neurospheres (Fig. 4c). The sSNV based clonal tracking plots for the paired patient samples identified two subclones in the HF3177 recurrence (Fig. 4d) that were not detected in the HF3016 neurosphere/PDX models, suggesting that these were independent of the *MYC* ecDNA element. Of note, a 0.5% cell frequency amplification was also detected in the parental tumor sample of HF2354, which increased to high levels in the derived neurosphere. DNA copy number analysis detected parallel *EGFR* and *CDK4* amplifications in the HF3016 primary GBM that were retained in HF3177 GBM recurrence as well as all model systems. Sequencing reads connecting the two amplifications and suggesting a complex structural variant were detected in the HF3016 neurosphere, the HF3016 PDXs, all HF3177 samples, but not the HF3016 primary GBM (Fig. 4e). Metaphase FISH on HF3016 neurosphere and HF3177 neurosphere confirmed that the *CDK4* and *EGFR* amplifications were part of the same ecDNA element (Fig. 4f). The genomic and extrachromosomal characteristics of these two tumor samples, their derived neurosphere cultures and xenografts provide an example of how multiple ecDNA elements are able to be preserved during tumor progression while in parallel acquiring new tumor subclones marked by sets of chromosomal sSNVs.

### Longitudinal maintenance of extrachromosomal DNA in patient tumors

Large, megabase sized double minutes are frequently found in glioblastoma and can be identified using whole genome sequencing and DNA copy number data ^21–23^. To determine whether extrachromosomal DNA can survive therapeutical barriers, we evaluated the DNA copy number profiles of 58 matching pairs of primary and recurrent glioma for the presence of ecDNAs ^4^. Evidence supporting the presence of ecDNA was found in 30 primary and 28 recurrent tumors spanning 34 patients and of these, ecDNA elements targeting cancer driver genes ^31^ were predicted in 22 primary tumors (Fig. 5a). The most frequently targeted gene was *EGFR* which was identified in 11 primary tumors, in agreement with previous reports^20,22^. *CDK4*, *PDGFRA* were detected in six and five primary tumors, respectively. We corroborated our computational predictions through interphase FISH analyses of 17 predicted ecDNAs and 26 non-altered loci across 6 primary/recurrent tumor pairs. Sixteen out of 17 genomic amplifications showed the highly variable number of DNA signals that is strongly suggestive of the extrachromosomal nature of the DNA locus (Fig. 5b, Supplementary Fig. 5a) whereas the 26 control DNA regions predicted to be non-amplified were confirmed as such (Supplementary Table 2). *EGFR* harboring ecDNA was preserved in the recurrent tumor in 4 out 5 pairs, half of which carried EGFRvIII mutation, including the HF2934 recurrent tumor analyzed after treatment with EGFR inhibitor dacomitinib (Fig. 5b, Supplementary Table 2). One tumor lost *EGFR* ecDNA and vIII mutation upon recurrence (HF2829), after treatment with the standard of care (radiation and temozolomide). In one case *MET* ecDNA was present in the primary tumor and maintained in the recurrence, while *MYC* ecDNA emerged upon recurrence, similar to what we reported above for the HF3016/HF3177 pair. To corroborate 55 DNA copy number predicted ecDNAs, we analysed whole genome and RNA sequencing data, which identified sequencing reads connecting adjacent focally amplified DNA segments (Fig. 5c and Supplementary Fig. 5b) supporting the predictions. After disease recurrence, 19 of 22 tumors preserved at least one cancer driver ecDNA, supporting the notion that ecDNA can prevail following the selective pressure imposed by anti-cancer therapy. We did not detect any significant correlations between somatic mutations and the presence of ecDNA. This analysis was potentially limited by the cohort size and our sensitivity in detecting ecDNA.

**Figure 5.**
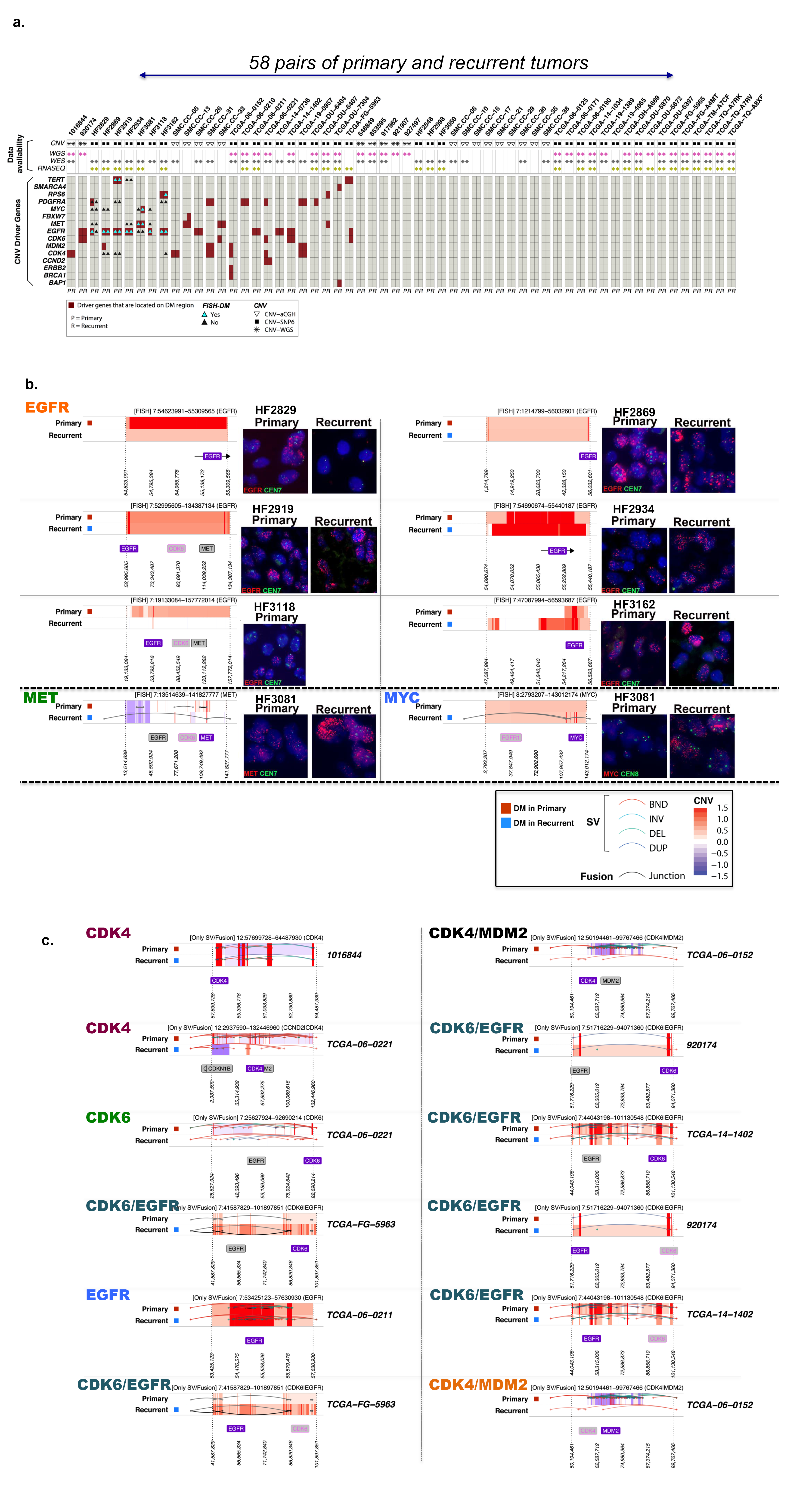
Copy number variant driver genes located on the potential double minute (DM) regions. **a.** 58 tumors (30 P, 28 R) from 34 patients were predicted to contain at least one ecDNA. Amongst these, 39 driver gene harboring ecDNAs were predicted in 22 primary tumors, of which 27 were also detected in the matching recurrent tumors. **b.** DNA copy number based predictions of double minute (DM) regions validated using fluorescent in situ hybridization in FFPE tissue sections. **c.** DNA copy number based predictions of double minute (DM) regions validated using whole genome or RNA sequencing.

## Discussion

Glioblastoma is a heterogeneous disease that is highly resistant to chemo- and radiotherapy. New modalities for treatment are urgently needed. Modeling of tumors through cell culture and orthotopic xenotransplantation are essential approaches for preclinical therapeutic target screening and validation, but in GBM have yet to result in novel treatments. To what extent these models truthfully recapitulate the parental tumor is a topic of active discussion. Here, we showed that neurosphere and orthotopic xenograft tumor models are genomically similar, capturing over 80% of all genomic alterations detected in the parental tumors.

EcDNA is increasingly recognized as playing an important role in tumorigenesis and gliomagenesis in particular ^20–23,30^. Our results provide direct evidence that ecDNA enhance genomic diversity during tumor evolution, and show how ecDNA elements can mark major clonal expansion in otherwise stable genomic background. Little is known about the mechanism through which these elements arise and how they become fixed across a cancer cell population. Our analysis provides a comprehensive study of the fate of chromosomal SNVs and ecDNA oncogene amplifications in GBM in a panel of tumors and derivative models. We further demonstrated the widespread presence of ecDNA driven oncogene amplification through extensive FISH analysis on sets of paired primary and recurrent tumor samples. Focal gene amplifications have traditionally been recognized as homogeneously staining regions (HSR) and these may originate from chromosomal insertions of ecDNA ^25^. We did not observe HSR-like staining patterns for the amplified genes in this study which suggests that this is not a common mechanism for gene amplification in GBM. We captured the early stages of *MYC* ecDNA expansion in the HF3016 and HF2354 tumors with 0.5-2% of cells presenting amplification (<30 copies/nucleus), with no evidence of chromosomal based gene amplification, while in all derived models, as well as the HF3016 recurrence (HF3077), the frequency of *MYC* amplification increased to 100% of cells with up to 100 copies/nucleus. These results are consistent with an origin through excision of a *MYC* containing chromosomal DNA segment and end-joining into a circular ecDNA, with subsequent amplification of the ecDNA^30^, followed by selection of *MYC*-amplified cells *in vitro* and in the recurrent tumor. Spindle assembly and chromosome segregation during mitosis lead to genetically identical daughter cells, containing similar sets of chromosomal sSNVs and DNA copy number alterations. Double minutes/ecDNAs are replicated during S-phase, but lack the centromeres that dictate the organization of the mitotic spindle, and as a result are randomly distributed across the daughter cells during mitosis. EcDNA elements thus inherit in a radically different fashion than chromosomes. This divergence in inheritance mechanism may explain for example why the evolution of the *MET* event was not similarly captured by sSNVs (Fig. 6), and shows that extrachromosomal elements play a key role in increasing genomic diversity during tumor evolution. Previous studies have found that extrachromosomal bodies can provide a reservoir for therapeutically targetable genomic alterations ^32^. Targeted MET inhibition of *MET* amplified GBMs has shown clinical promise ^33^, although the variable responses to MET inhibition recorded in our data suggest that single MET inhibiting agent efficacy is influenced by other factors. Our observations extend recent findings that ecDNA are frequently detected in cancer ^20,22^ and demonstrate that detection of point mutations alone is insufficient to accurately delineate tumor evolutionary process. The disappearance of extrachromosomally amplified driver genes in neurosphere cultures has been reported ^23^, but is not confirmed in our our data that show ecDNA carrying amplification of *MYC*, *CDK4*, *EGFR*, and *PDGFRA* were maintained in neurosphere cultures, at least up to passage 18.

**Figure 6.**
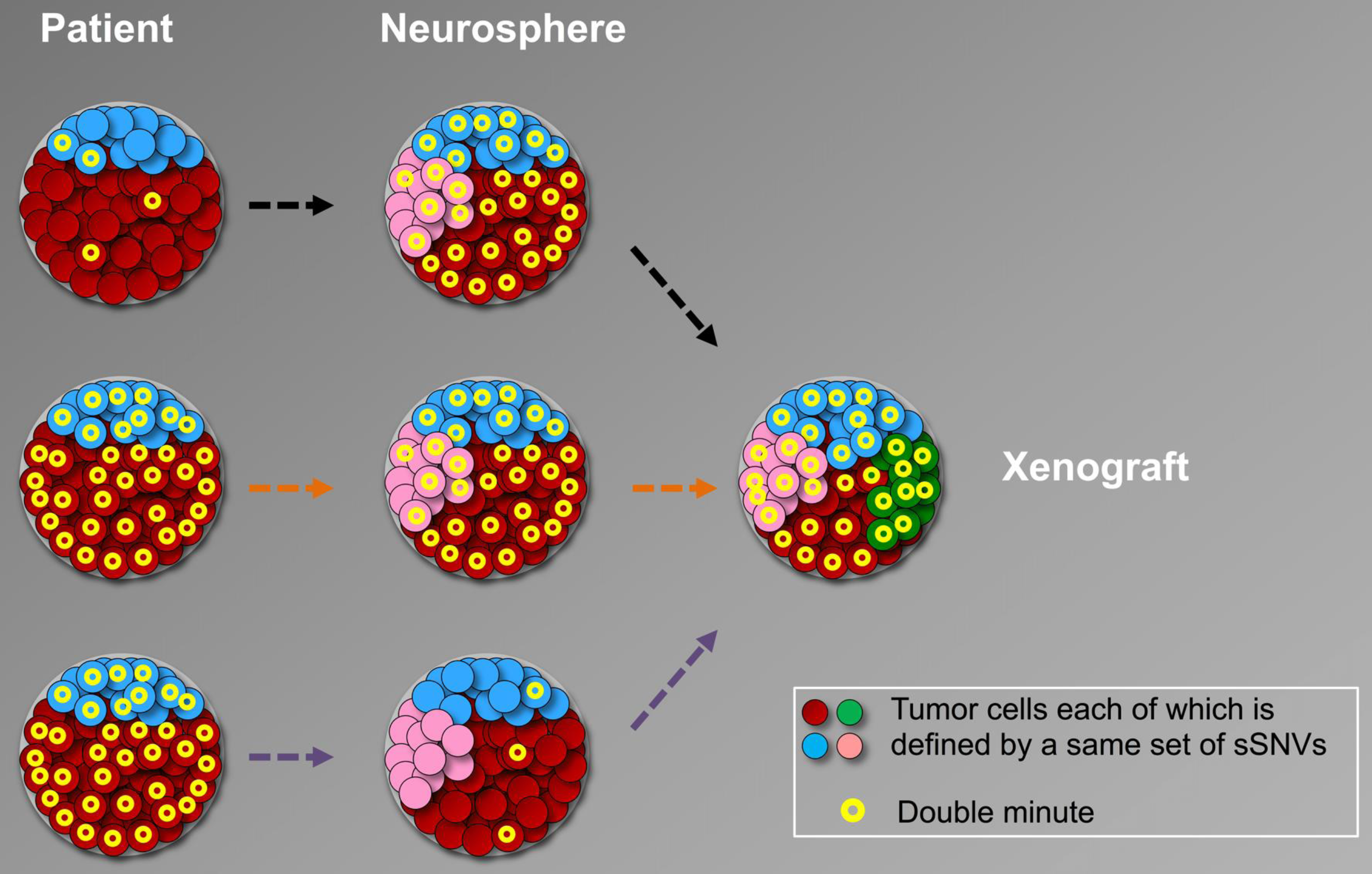
Schematic illustration of double minute contribution to clonal evolution in GBM patient derived models. The proliferation patterns in GBM tumors and models in which double minutes provide the dominant evolutionary force.

Double minutes have been reported in 10-40% of GBM ^21–23^. These lesions frequently involved genes on chromosome 12p, including *CDK4* and *MDM2*, span up to several megabases in size, and can be recognized by an intermittent amplification-deletion DNA copy number pattern. An important contribution of our work is in the size of the ecDNA elements identified, which ranged from several kb to several Mb. Larger ecDNAs are often characterized by an intermittent amplification-deletion DNA copy number pattern^22^, but kb-sized single segment episomes can only be identified using high throughput sequencing approaches^34^ or DNA staining/FISH experiments ^20^ and are therefore likely underreported. Whether ecDNA size and structure affects the mechanism of tumorigenesis is unclear. Extrachromosomal DNA is an understudied domain in cancer. Our analysis emphasizes the importance of this genomic alteration category for gliomagenesis. Future studies that specifically target the formation of episomal events may lead to therapies to prevent this process from happening. The models we described here may play a pivotal role in evaluating the potential of such approaches.

## Acknowledgments

The authors would like to thank Dr. Norman Lehman and Dr. Chunhai (Charlie) Hao for pathology reviews; Lisa Scarpace for clinical information; Susan Irtenkauf, Laura Hasselbach, Kevin Nelson, Kimberly Bergman, and Susan Sobiechowski for cell culture and animal work; Andrea Transou, Yuling Meng, and Enoch Carlton for histology at HFH. We thank Genevieve Geneau, Sharen Roland, and Pac Bio platform personnel of the Génome Québec/Genome Canada-funded Innovation Centre for providing Pacific Biosciences sequencing. This work was supported by the LIGHT Research Program at the Hermelin Brain Tumor Center; grants from the National Institutes of Health P50 CA127001, R01 CA190121, P01 CA085878 and P30CA034196; the Cancer Prevention & Research Institute of Texas (CPRIT) R140606. This work was also supported by a grant of the Korea Health Technology R&D project through the Korea Health Industry Development Institute (KHIDI), funded by the Ministry of Health & Welfare, Republic of Korea (HI14C3418) We are hugely indebted to the patients who provided tumor and germline material for the purpose of this study.

## Author Information

BAM files from exome sequencing, low pass whole genome sequencing and RNA sequencing used in this study were deposited to the European Genome-phenome Archive (EGA; http://www.ebi.ac.uk/ega/), which is hosted by the EBI and the CRG, under accession number EGAS00001001878. The authors declare no competing financial interests.

## Supplementary Information

Supplementary Figures, Methods, and Supplementary Tables are available as supplementary data.

